# Genetic diversity and phylogeography of swine influenza A virus in 2013-2022 in Europe using a harmonized genotyping nomenclature

**DOI:** 10.64898/2026.09.21.753142

**Authors:** Gautier Richard, Benjamin C. Mollett, Pia Ryt-Hansen, Séverine Hervé, Martí Cortey Marquès, Enric Mateu, Ana Moreno, Chiara Chiapponi, Lars Erik Larsen, Timm Harder, Helen E. Everett, Gaëlle Simon

## Abstract

Swine Influenza A virus (swIAV) poses constant health threat for both humans and animals. This study comprehensively analysed the genetic diversity and evolution of European swIAVs from 2013 to 2022. To address existing classification gaps, an easily accessible full-genome genotyping tool based on Nextclade was developed alongside a global nomenclature system. Application of this tool to European sequence data revealed unprecedented genetic diversity, successfully identifying 150 swIAV genotypes mixing segments of both swine and human origin. Bayesian phylogeographic models, combined with economic data, identified that the trade of live weaner pigs for fattening drove the transboundary spread of swIAVs across Europe. The study detected repeated human-to-swine reverse-zoonotic events across multiple European countries. These findings clearly highlight the role of commercial pig movements in viral dissemination and emphasize the critical importance of harmonized genomic surveillance to monitor swIAV viral evolution to inform global pre-pandemic preparedness strategies.

## Introduction

Animal Influenza A viruses (IAVs) pose a constant threat to human and animal health. Their broad host range and segmented RNA genome enable rapid genetic diversification through the accumulation of mutations (drift) and infrequent gene segment exchange (shift) ^1^. Domestic pigs serve as crucial host metapopulations for viral maintenance and diversification since they are susceptible to IAVs originating from diverse mammalian (especially human) and avian species. They function as mixing vessels for emergence of novel reassortants strains with unique gene constellations and pandemic potential. Given the high risk of interspecies spillover at the human-animal interface and owing to the swine-origin 2009 H1N1 pandemic, swine IAV (swIAV) genetic diversity is monitored by the swine influenza technical group of OFFLU, a joint World Organization for Animal Health (WOAH) & Food Administration Organization (FAO) network for swine influenza. IAV antigenic properties depend on the envelope glycoproteins Hemagglutinin (HA) and Neuraminidase (NA), which OFFLU classifies using a global nomenclature system ^2^, and report HA/NA diversity biannually to the WHO vaccine candidate meeting, enabling rapid provision of candidate strains if vaccine composition needs urgent updating.

SwIAV was first detected in the early 20^th^ century in Europe and North America, termed classical swine H1N1 (1A.1) lineage, sharing a common ancestor with the 1918 pandemic influenza ^3–9^. This lineage circulated in Europe until supplanted by a novel “Eurasian avian-like” (“av” or “EA”) H1N1 lineage (H1C/N1EA), first detected in 1979 following an avian-to-swine transmission event ^10^. The H1C lineage remained enzootic in European herds, continuing to reassort and evolve ^11–15^.

Subsequent reverse-zoonotic spillover of human viruses into pigs caused two major genetic shifts in Europe. First, a 1968 human-origin ‘Hong Kong’ H3N2 virus entered European herds in the early 1970s ^16–18^ and acquired the EA internal gene cassette from H1C viruses, forming the “reassortant swine H3N2” lineage H3.1970.1/N2G (N2 Gent/1984) which co-circulated in the 1980s with the enzootic H1C viruses ^2,19–21^. Second, in 1994 in the United Kingdom, this swine H3N2 lineage acquired a HA from a human-origin H1N1 virus (A/Chile/1983-like strain) and formed the “human-like H1N2” (H1B.1 and “N2 Scotland/1994”, referred to as N2S) virus. This lineage subsequently became established in several countries in continental Europe ^2,22,23^. Further adaptation and reassortment in pig herds then yielded the H1B.1.2.2 lineage in Italy ^15^ and later in Europe, which is characterized by the HA 146-147 amino-acids deletion and by an NA derived from a 1998 human seasonal H3N2 strain (N2EU) ^24^. A further reassortant, “H1N2dk,” was identified in Denmark in 2003, combining H1C-lineage segments with an N2G segment from H3.1970.1 ^25^, and subsequently spread across Europe ^11,14,15,26,27^. Meanwhile, the prevalence of the original H3.1970.1/N2G virus has steadily declined across Europe since the mid-1990s ^28^.

In April 2009, an H1N1 pandemic emerged following zoonotic transmission of a swIAV combining gene segments from American swIAV lineages with NA and matrix (M) segments from the European H1C lineage — a reassortant not previously identified in pigs^2,29^. A putative precursor was later found in Mexican pig herds ^30^. A/H1N1pdm09 likely infected European pigs in September 2009, causing respiratory disease in growing and finishing pigs ^31,32^. It was subsequently detected in most pig populations in Europe and globally ^14,15,33–35^ ^13,36–38^, and designated as the H1A.3.3.2 clade. Having replaced the previous seasonal H1N1 in humans ^39^, A/H1N1pdm09 has since been a recurring source of human-to-swine spillover ^40^, substantially increasing swIAV diversity and yielding additional novel genotypes ^11,13–15,37,38,41,42^.

Coordinated European efforts such as ESNIP1-3 ^14,15^ and a limited 2015-2018 study ^11^ have characterised swIAV diversity, but the mechanisms and phylogeographic patterns of swIAV spread remain poorly understood. To address this gap, we describe evolutionary changes in European swIAVs over 2013-2022 using whole-genome sequence data from six major pork-producing countries, revealing continuing genetic diversification. Since an accessible, full-genome genotyping tool with an index-independent nomenclature was needed, we propose an updated European genotyping system harmonized with the global phylogeny-based framework ^2^ enhanced by an automated genotyping tool. Finally, we employed phylogeographic analyses to characterise swIAV transmission dynamics and reverse-zoonoses across Europe.

## Results

### A harmonized international tool to genotype swIAVs

Given the genetic diversity of swIAV in Europe and data gaps since 2013, an appropriate phylogenetic reference dataset was assembled, annotated, and made available as a tool to enable fast and accurate swIAV genotyping. Ten Nextclade builds based on contemporary H1, H3, N1, N2, PB2, PB1, PA, NP, M, and NS phylogenies were generated (Figure 1A) using an annotated international swIAV dataset based on OctoFlu ^2,43^. These builds were made available as a web interface (https://clades.nextstrain.org/) and are compatible with a new command-line tool for swIAV full-genome batch genotyping (https://github.com/gtrichard/influenza_sequences_toolbox/blob/main/bin/nextcladeGenotype).

**Figure 1.**
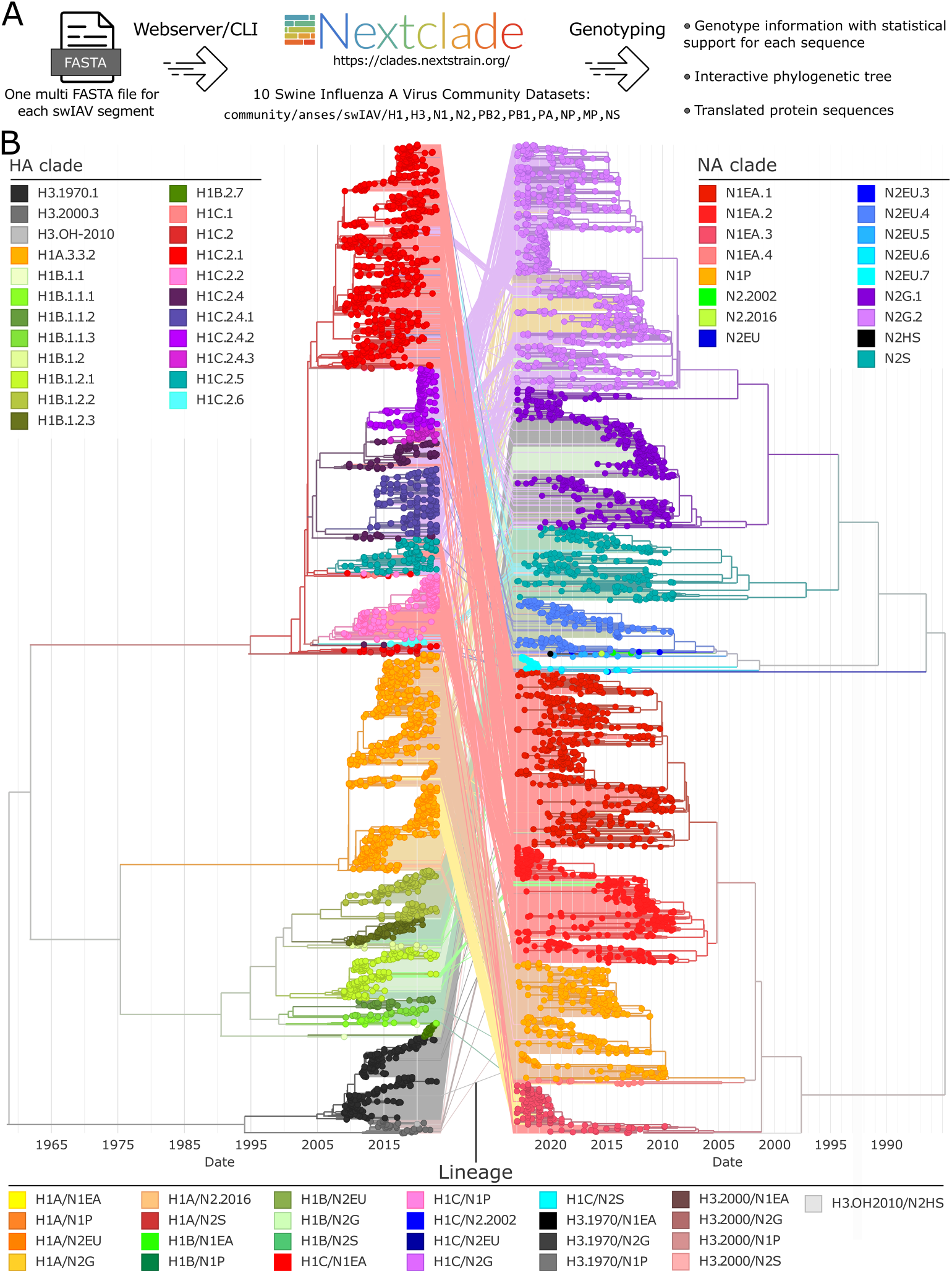
Harmonised genotyping tool for all swIAV segments and the resulting classification of HA and NA segments on 2009-2022 European swIAV sequences. **A)** Workflow of usage of the novel genotyping tool for swIAVs based on Nextclade, which can be used as both a command-line tool and a web-based tool to genotype sequences contained in an unaligned fasta file. The tool consists of 10 independent Nextclade builds (H1, H3, N1, N2, PB2, PB1, PA, NP, M, NS) maintained at https://github.com/nextstrain/nextclade_data/tree/release/data/community/anses/swIAV and hosted at https://clades.nextstrain.org/. **B)** Maximum-likelihood HA and NA tanglegram. HA sequences (left) are coloured based on the HA clades while NA sequences (right) are coloured based on the NA clades. Lines (middle) connect the HA and NA sequences for each strain and are coloured based on a “lineage” nomenclature capturing the higher-level lineages of HA and NA classifications. Interactive phylogenies at: https://nextstrain.org/community/gtrichard/swIAV-Europe-2013-2022.

Based on the OctoFlu database and the proposed swIAV global phylogeny clade annotation rules, four new sub-clades in the N1 Eurasian avian-like lineage (N1EA.1-4) and a new N2EU.7 clade were identified (Supplementary Table S1). Internal gene segments were classified at a coarser level than HA/NA, as Eurasian avian-like (EA), Pandemic 2009 (pdm), or Human Seasonal (HS, corresponding to recent (>2020) H3N2 reverse zoonoses) to remain informative at the level of Europe.

This tool was applied to swIAV sequences sampled in Europe from 2009-2022 to characterize their diversity between 2013 and 2022, i.e. after the conclusion of ESNIP3 ^15^, and to evaluate the impact of H1A.3.3.2 viruses introduced after 2009 (Figure 1B). The reference dataset included all publicly available European swIAV sequences and metadata from GISAID EpiFlu and NCBI Virus, supplemented by previously unreleased data from six partner countries (Denmark, France, Germany, Italy, Spain, and the UK), made available through NCBI (Supplementary Table S2). Classification of the HA and NA segments (Figure 1B) showed no apparent mislabelled branches while providing more fine-grained diversity classification for both segments. No mislabelling was evident for the remaining segments either.

### Diversity of swIAV in Europe between 2013 and 2022

Based on the classification of swIAV HA and NA, different levels of diversity were determined, with the higher level order being “subtype” corresponding to the HA and NA numbering (i.e., H1N1); followed by “lineages” corresponding to the first level of the HA and NA clade definition (i.e., H1C/N1EA); and finally the “HA/NA clades”, corresponding to the combined complete clades (i.e., H1C.2.1/N1EA.1) detailed in Supplemental Table S1. The most prevalent lineages detected in Europe between 2013 and 2022 were the H1C/N1EA found in 14 countries and predominant in 12 of them, followed by the H1A/N1P found in 12 countries but only predominant in 5 of them (Figure 2A and 2B). According to the HA/NA tanglegram (Figure 1B), the most common HA/NA pairings were H1C.2.1/N1EA.1, H1C.2.2/N1EA.2 and H1A.3.3.2/N1P. Interestingly, H1C/N2G was the third most widespread lineage identified in 9 countries, including Denmark where it was the predominant lineage, and France and Spain where this lineage was almost as prevalent as H1C/N1EA. The underlying HA/NA clades of the H1C/N1EA lineage displayed a high diversity compared to other lineages, with 17 underlying HA/NA clades combinations compared to 15 combinations of the H1C/N2G lineage (Figure 3). This analysis highlights the ongoing diversification of the H1C.2.4 strains in Europe, with 3 subclades and their association with 10 NA clades. Despite the H3N2 progressive disappearance in Europe among the swine population, with only 12 strains in 2021-2022 compared to 69 strains in 2013-2014, the H3.1970/N2G remained the fourth most widespread lineage in Europe between 2013 and 2022 (found in 8 countries, at low prevalence, Figure 2). These strains were continuously reported in Italy (Figure 3) and detected between 2013 and 2017 in France, Germany and Spain.

**Figure 2.**
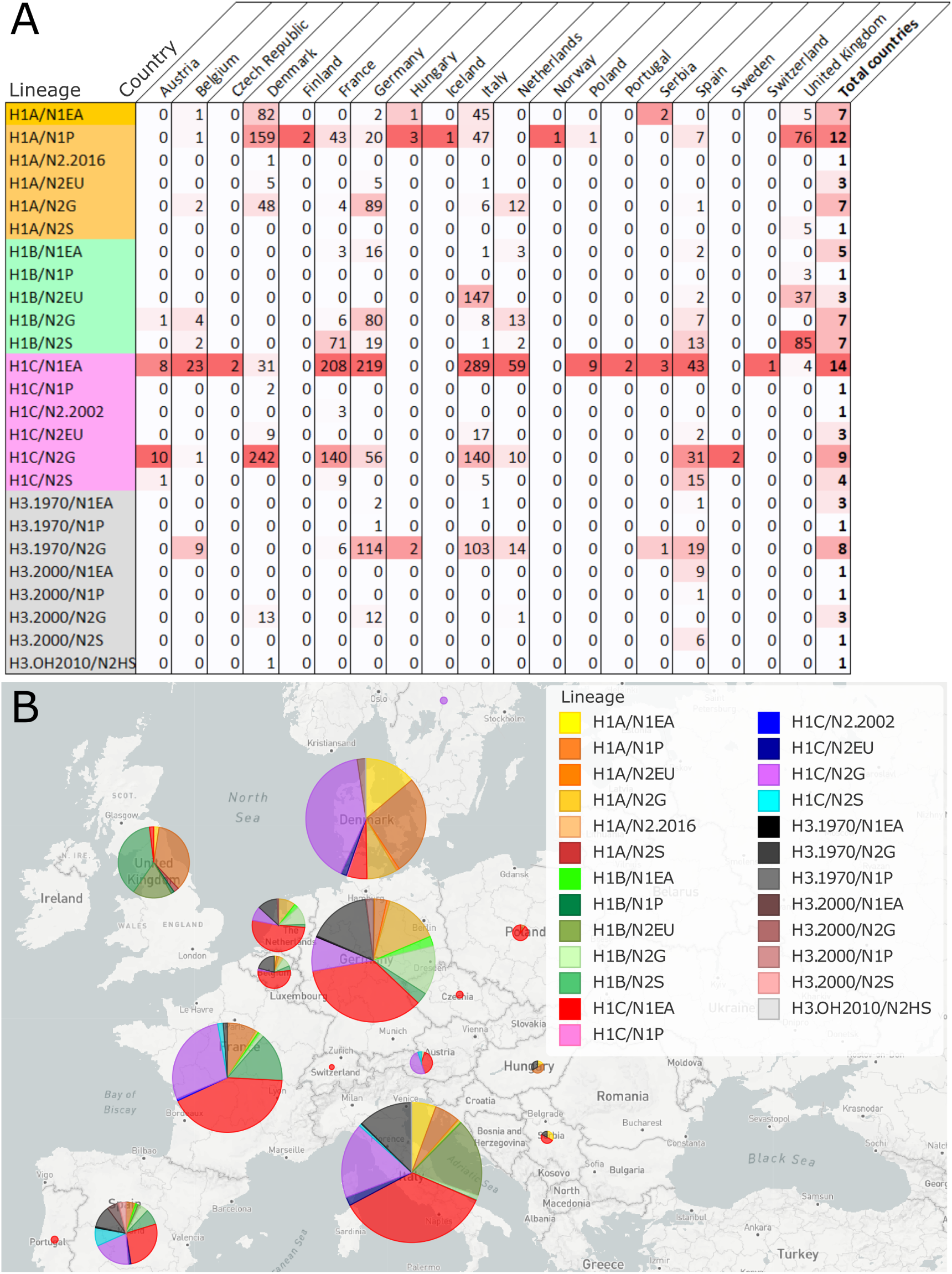
SwIAV Lineages distribution in Europe between 2013 and 2022. **A)** Number of strains sequenced per subtypes per country. The table is coloured based on the representativity of each lineage within each country. The last column represents the number of countries in which each lineage has been detected for the considered period. **B)** Map of proportion of each lineage per country based on the sequenced strains per country.

**Figure 3.**
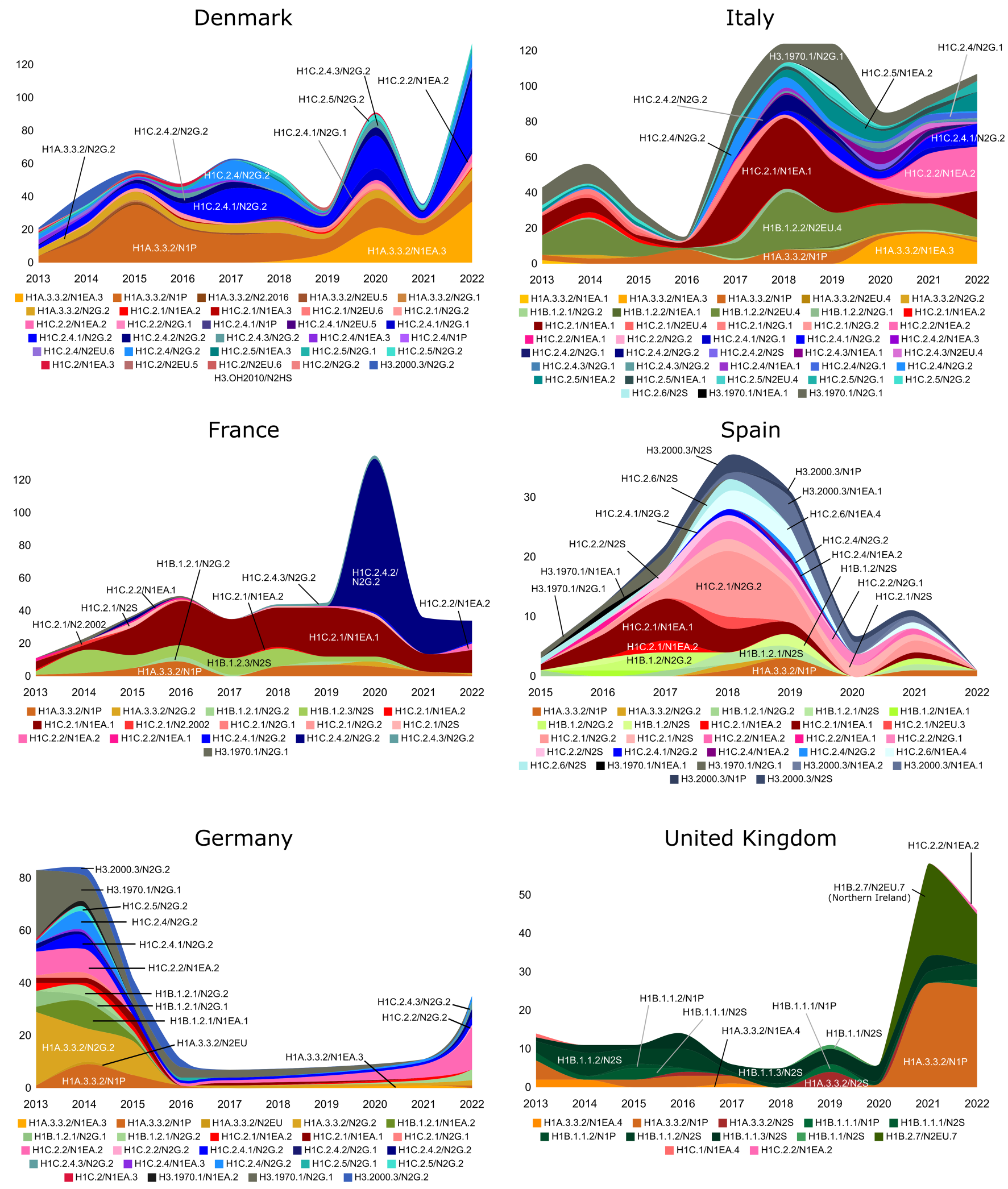
Distribution of swIAV combined clades annually between 2013 and 2022 in six European countries. Number of genotyped viruses are represented on the Y-axis per year (on the X-axis). The combined clades detected in each country, following the HA-clade/NA-clade nomenclature, are represented below each streamgraph with their respective unique colours.

Novel swIAV reassortants were also detected in several countries between 2013 and 2022, including H1C.2.4.2/N2G.2 in France (from 2020-), H1A.3.3.2/N1EA.3 in Italy and Denmark (2019-), H1C.2.2/N1EA.1 in Italy (2020-), and H3.2000.3/N2G.2 in Denmark and Germany (2014-). New clades were also identified in Europe within this period compared to previous European studies, such as H1B.2.7 and N2EU.7 clades in Northern Ireland from 2021 onward, the H1C.2.4 and H1C.2.5 clades throughout Europe, which were defined in February 2021 in the OFFLU Swine Influenza VCM (https://offlu.org/technical-activities/february-2021-swine-influenza/), and the H1C.2.6 clade identified in Spain from 2015 onward.

### Full-genome swIAV sequence analyses reveal high genotype diversity and complex reassortment patterns within Europe

The swIAV genotype was assigned using the classification tool that employed the full HA clade, the full NA clade, followed by the first letter of the clade for the remaining gene segments, ordered according to convention as PB2, PB1, PA, NP, M, and NS, which can be of clades P = H1N1pdm09 lineage, E = Eurasian avian like or H = H3 human seasonal. This gave for instance H1A.3.3.2/N1P/PPPPPP for the gene constellation associated with the H1N1 swIAV derived from the A/H1N1pdm09 virus. Such a nomenclature was directly obtained from the newly developed nextcladeGenotype tool.

This analysis permitted the identification of 150 swIAV genotypes in nine countries with enough sequences between 2013 and 2022 (Figure 4). Most genotypes were found to be country-specific and sporadic. The most widespread genotypes retained non-reassortant gene constellations and were H1C.2.2/N1EA.2/EEEEEE (eight countries), H1C.2.1/N1EA.1/EEEEEE, H1A.3.3.2/N1P/PPPPPP, H1A.3.3.2/N2G.2/PPPPPP and H1B.1.2.1/N2G.2/EEEEEE (seven countries each), as well as the H1C.2.1/N1EA.2/EEEEEE which was found in eight countries, but remained sporadic in most. Some reassortants of the internal genes were widespread in Europe for the considered period, such as H1C.2.2/N1EA.2/EEEEPE (five countries), H1C.2.4/N2G.2/PPPPPE (four countries) and H1A.3.3.2/N1EA.3/PPPPPE (three countries). Most single gene reassortment affected the M (26 genotypes) and NS (20 genotypes) genes compared with PB2 (2), PB1 (5), PA (2), and NP (4) genes. While many genotypes were found in single countries, others were reported in two countries or more, with for instance four genotypes that were only found in Denmark and Italy, raising the question of potential swIAV co-evolution or transmissions between countries.

**Figure 4.**
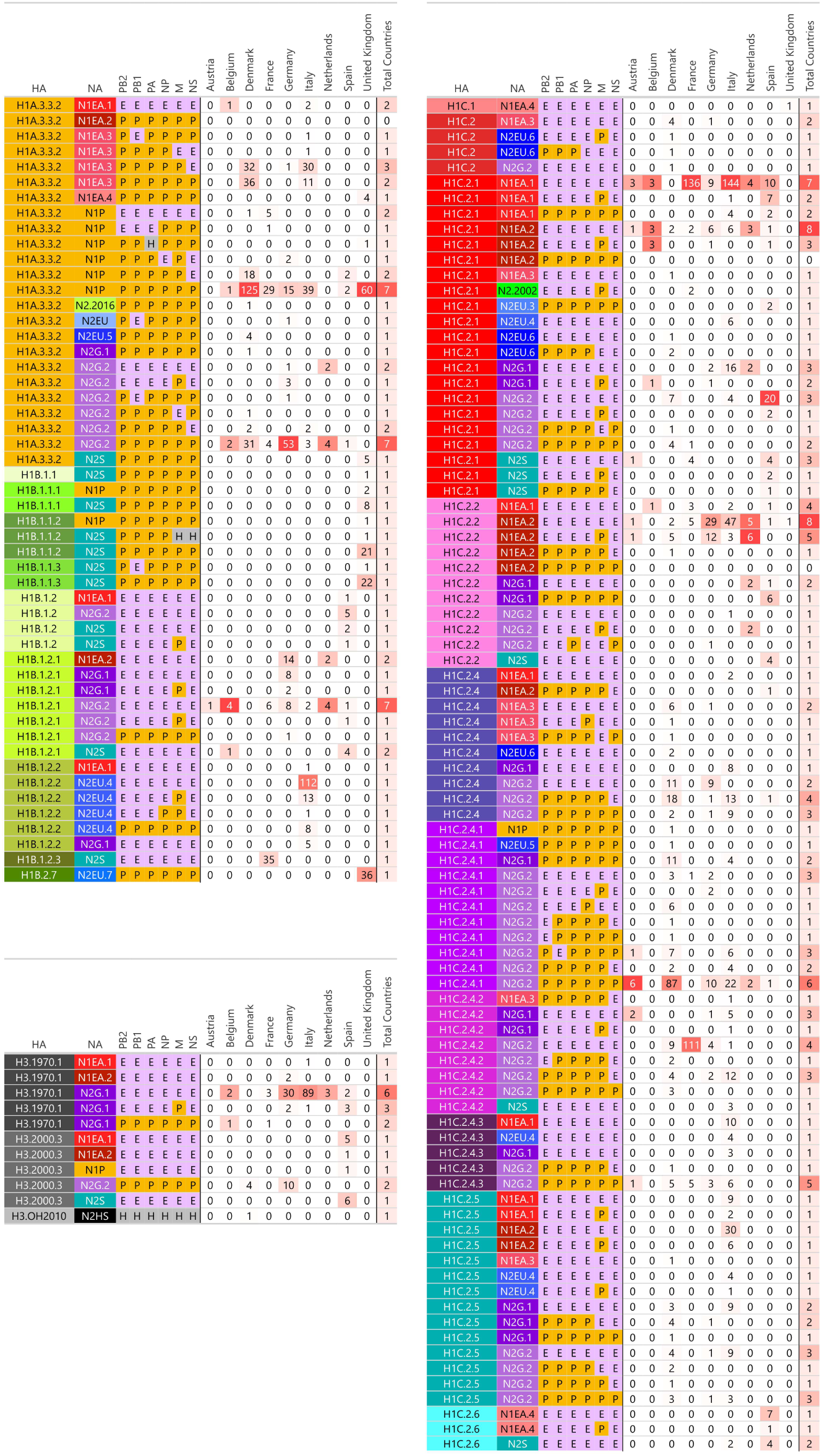
SwIAV genotypes per country with 10 or more full genome sequences characterised from samples collected between 2013 and 2022. Genotypes were represented using a harmonised nomenclature such as HA-clade/NA-clade/GGGGGG where G represents the first letter of the clade attributed to each internal gene in the numerical order (PB2, PB1, PA, NP, M, NS) with E = Eurasian avian-like, H = human seasonal, P = pdm09. The numbers are coloured depending on the number of each genotype within a given country.

### Inter European transmission patterns of H1C swIAVs

To investigate viral dissemination between European countries, a phylogeographic Bayesian approach using BEAST X was conducted on the HA 1C phylogeny, the most widespread HA lineage. Sequences were filtered to include only countries with at least 10 sequences reported between 2013 and 2022, producing a Maximum Clade Credibility Tree (Figure 5A) after robust computations (Supplementary Table S1). Markov Jumps (MJ, Figure 5B), representing the frequency of swIAV transmissions between countries along the computed trees, and Bayes Factors (Figure 5C), representing statistical support for MJ, were calculated, identifying many recurrent and statistically supported swIAV transfers between European countries. Of the eight analysed countries, strains were identified to disseminate primarily from Germany to other countries followed by transmission of viruses from The Netherlands and Denmark to five countries. The most frequent transmissions were from Denmark to Germany (33.87 MJ), Denmark to Italy (31.42 MJ), and Germany to the Netherlands (23.87 MJ). A Maximum-Likelihood-based mugration test, where arc width represents transmission frequency between countries, showed similar patterns (Figure 5D).

**Figure 5.**
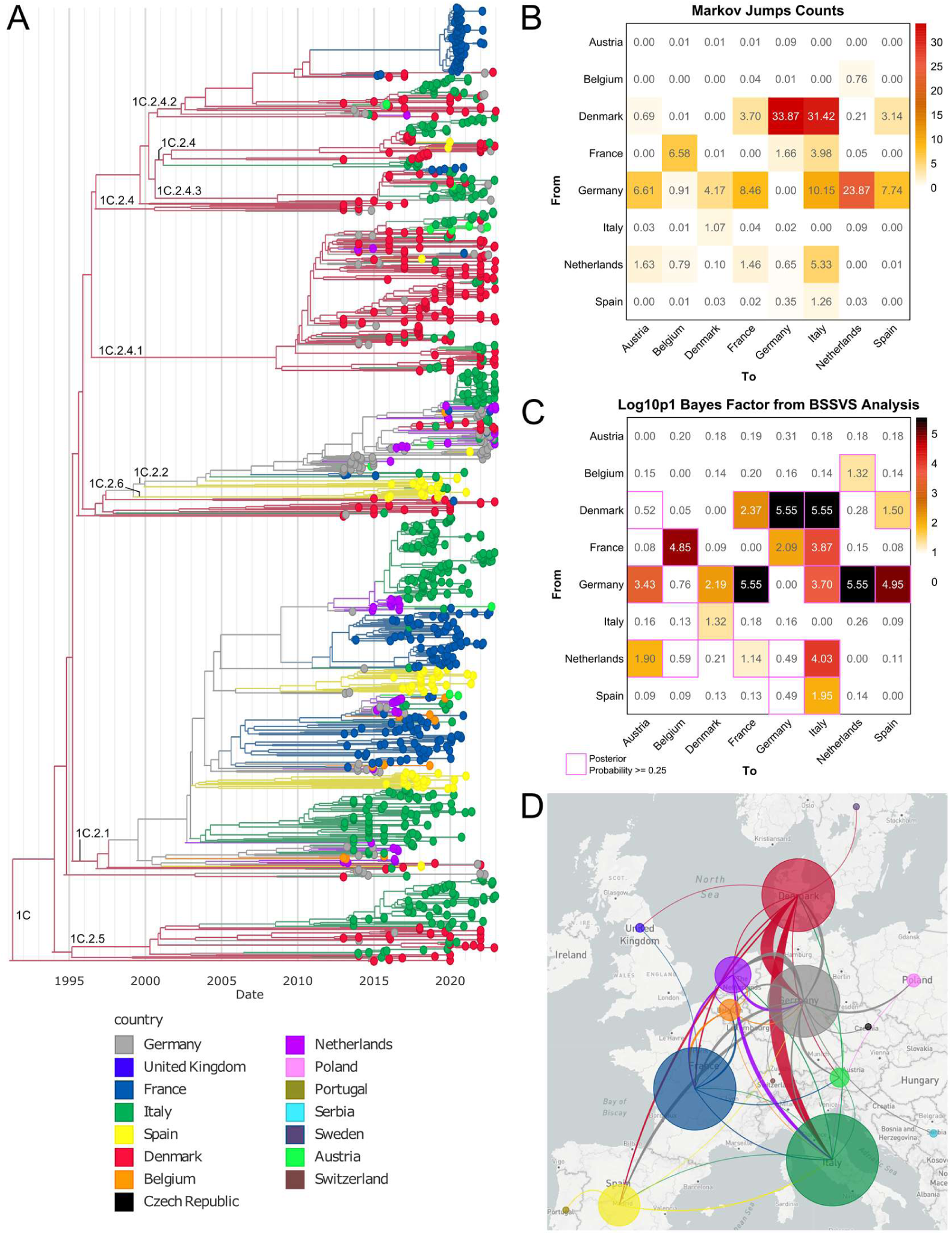
Spillovers from country to country of the most widespread HA lineage in Europe between 2013 and 2022: HA 1C. **A.** BEAST Maximum Clade Credibility (MCC) tree based on HA 1C sequences from country with at least 10 strains sequences (Austria, Belgium, Denmark, France, Germany, Italy, The Netherlands and Spain), coloured by country. **B.** Markov Jump Counts calculated along the BEAST trees which were used to build the BEAST MCC. Represents the swIAVs spillovers frequency between countries with directionality (from on the Y-axis, to on the X-axis). **C.** Log10+1 of the Bayes Factors extracted from BSVSS BEAST analysis. Represents the statistical support of the Markov Jumps. **D.** Maximum-likelihood mugration test calculated with TreeTime on the HA 1C phylogeny, representing swIAVs spillovers between countries. The larger the links, the higher the number of sequences involved in spillovers between two countries. The links colour is based on the country of origin of the swIAVs.

To identify potential drivers of swIAV transmissions between European countries, H1C phylogenetic data were combined with trade volumes of live swine weighing under 50 kg (mainly growing pigs, 7-30kg, corresponding to weaning pigs, hereafter “weaners”) and live breeding swine (”breeders”) using a BEAST GLM approach. Swine over 50 kg (excluding breeders, mainly slaughter pigs) were excluded, since these are typically traded for immediate slaughter and do not move farm-to-farm between countries ^44^. Average annual gross exports (USD) of these two live-pig types were calculated per country over 2013-2022 using the WITS database (Figure 6). Export destinations and volume proportions per country (second ring, Figure 6) differed between the two types, indicating distinct exchange routes and volumes for weaners versus breeders within Europe. Integrating H1C phylogeny data with weaners and breeder’s gross exports data revealed that at least one trade matrix was used consistently across the Markov Chain (country.includePredictor=1.066, Supplementary Table S4), consisting almost exclusively of the weaners trade matrix (country.coefIndicators_weaners=1, stdev=0 and country.coefIndicators_breeders=0.007, stdev=0.25, Figure 5C-D). Moreover, swIAV transmissions between European countries were strongly associated with weaners trade (country.coefficients_weaners=1.689, stdev=0.17, Figure 5D), while breeders trade had limited impact (country.coefficients_breeders=0.0003, stdev=1.92 Figure 5C). Taken together, this suggested active swIAV transmissions between European countries through trade of infected live weaning pigs.

**Figure 6.**
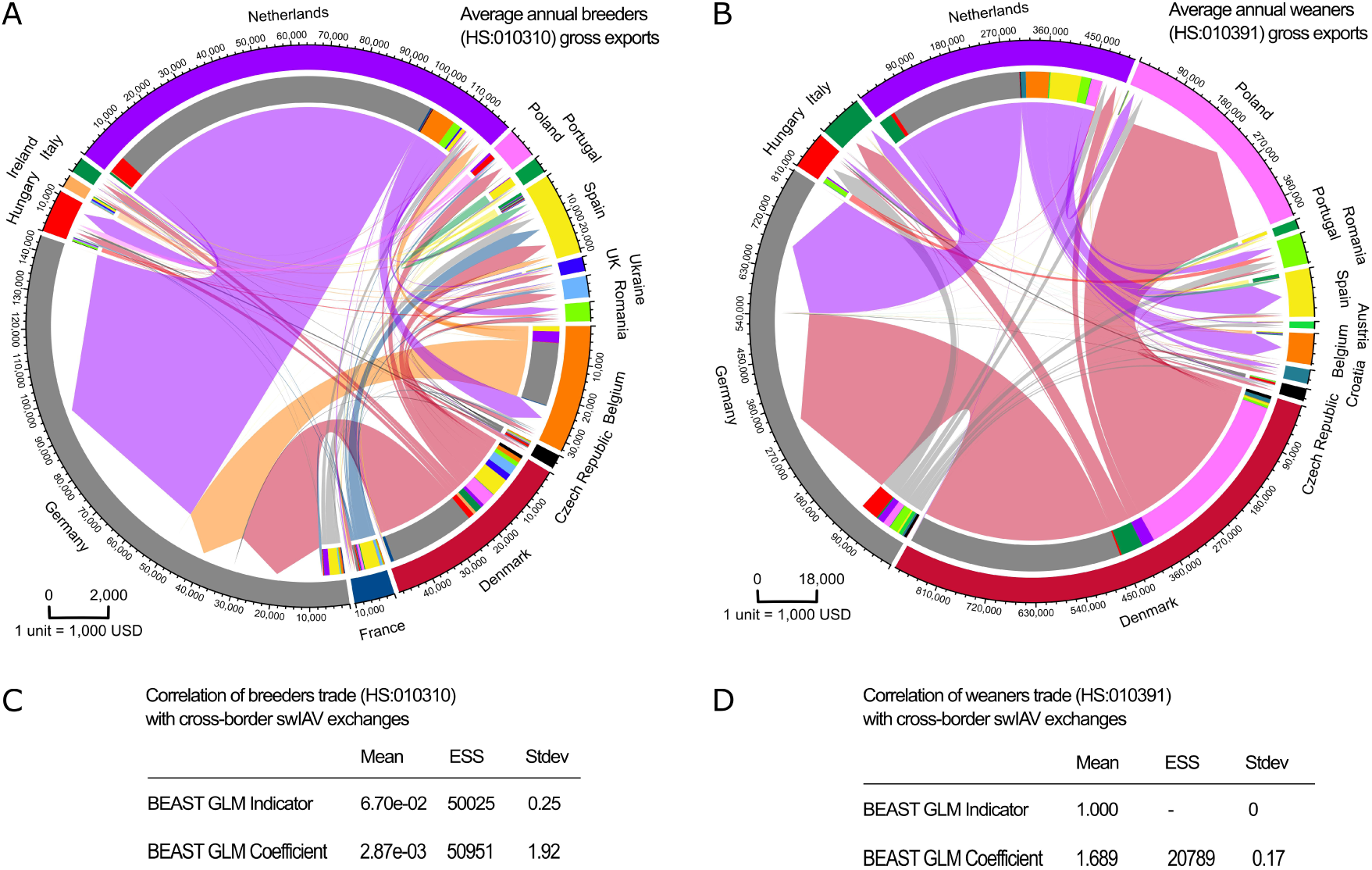
Average annual live-swine trades in Europe between 2013 and 2022. **A and B:** Average annual gross exports (trade volumes) expressed in 1,000 USD for live-swine of the pure-bred breeder type (HS:010310, panel A, “breeders”) or weighting less than 50 kilograms type (HS:010391, panel B, “weaners”) from the World Integrated Trade Solution database. Outer circle: each country has a colour and the radius of the country corresponds to gross exports expressed in 1,000 USD. For each panel, the unit of one tick is represented in the bottom left corner: 1 tick corresponds to 2,000 units of 1,000 USD for panel A and 18,000 units of 1,000 USD for panel B. Inner circle: gross exports volume (radius) and destination (colour). Central arrows: trade flows from one country to another with a colour based on the country of origin and a size proportional to the gross export volume. **C and D:** BEAST GLM Predictors statistics, combined based on two runs, of HS:010310 (panel C, breeders) and HS:010391 (panel D, weaners) trade values between countries in relation with the H1C swIAV phylogeny and exchanges between countries described in Figure 5. Indicator determines how often the live-swine trade matrix was used alongside the Markov Chain to predict swIAV exchanges between countries (0 = never used, 1 = always used). Coefficient determines how strongly the live-swine trade values impact the swIAV exchanges between countries (<0.5 weak impact, >1 strong impact). Detailed statistics of the BEAST runs are available in Supplementary Table S4. ESS: Effective Sample Size. Stdev: Standard deviation.

### IAV reverse-zoonoses in Europe

As previously mentioned, H3N2 viruses detections were low between 2013 and 2022 in the swine population in Europe, especially concerning human seasonal H3 clades (2010 and 2020) since only a single H3.Other-Human.2010 sequence was reported in Denmark. Regarding human-seasonal H1N1 viruses and considering the genetic proximity with swine H1A.3.3.2 clade, the regularly updated H1 Nextclade human seasonal reference set using the A/California/07/2009 strain as a reference was used on swIAV H1A.3.3.2 sequences to assign a probability of being a reverse-zoonosis (Supplementary Figure S2). H1A.3.3.2 swIAV strains that were closely matching human seasonal H1 sequences according to Nextclade Quality Control score (reported as “High” in Supplementary Figure S2) were found in Belgium, Denmark, France, Germany, Italy, Spain and the United Kingdom. None were found as potential reverse-zoonosis in The Netherlands sequence dataset. The proportion of H1A.3.3.2 cases associated with a high probability of reverse zoonosis was found to be of 21% in average over five countries with enough data; with a high proportion of 60% in France.

## Discussion

SwIAV surveillance and genetic characterization is critical for animal welfare, pandemic preparedness and vaccine design ^45^, as exemplified by the 2009 swine-origin H1N1 pandemic. SwIAVs circulate globally and often become endemic in intensive farming systems ^46,47^, promoting persistence, co-infection and reassortment ^48^, leading to new viral genotypes with pandemic potential. SwIAV diversity was analysed at the scale of Europe by the ESNIP3 program until 2013, which showed a predominance of H1C/N1EA strains in Europe (53.6%) followed by H1N2, H3N2 and H1N1 pandemic lineages represented in almost equal proportions, with differences of incidence in each country ^14^.

Since HA and NA classifications evolved since 2013, and that it was difficult to assign swIAV genotypes in a simple and harmonized manner, a tool was developed in the present study based on Nextclade to assign clades to the eight segments of swIAV, accompanied by a nomenclature to describe swIAV genotypes detected in Europe. The proposed swIAV Nextclade tool is available as both a command-line tool and a web tool, making swIAV genotyping available to a broader audience compared to command-line specific genotyping tools such as OctoFlu ^43^. The present tool is also more accessible than the BV-BRC ^49^ classification web tool, which requires a login, the upload of viral sequences and the management of a file system. Since Nextclade web and command-line versions run both on the user’s machine locally, no sequence data are stored in a remote server, ensuring data privacy. Compared to the previously published swIAV genotype nomenclature, the newly proposed nomenclature in the form of HA/NA/GGGGGG (with G, the first letter of internal gene origin in numerical order) allows an easy update of the swIAV segment classification without the need to regularly update an index. It therefore has the capacity to accommodate the definition of new clades or subclades for any segment, compared to previous tools and nomenclatures ^11^. Such a system will also enhance comparisons between swIAV studies by allowing a rapid assessment of swIAV diversity in a unified manner, by avoiding confusion between different nomenclature systems, and by revealing novel insights including new reassortants or the need to define new clades based on phylogenetic analyses.

The application of the Nextclade swIAV genotyping tool to the swIAV positive samples collected in Europe between 2013 to 2022 revealed 150 genotypes, which has increased considerably since previous reports ^11,15^. This is due to a longer period being covered in the present study (10 years compared to 3-4 years), to the availability of more samples, and to the more detailed classification of HA and NA sub-clades. This will allow an enhanced comparison of swIAV surveillance between European countries and underpin informed decisions on future antigenic cartography studies and vaccine design.

Analyses of swIAV diversity between 2013 and 2022 revealed that viruses bearing H1A.3.3.2, H1C.2.2 or H1C.2.1 were the most common and widespread in Europe, consistent with previous studies ^11,15^. During the studied period, some countries reported emergence of new genotypes such as H1C.2.2/N1EA.2 in Italy, H1C.2.4.2/N2G.2 in France and H1A.3.3.2/N1EA.3 in Denmark and Italy ^50–52^. Within the H1C lineage, our analyses revealed an on-going diversification of the swIAV H1C.2.4 clade in Europe as well as an increase in detection overtime since 2013 in Denmark, Italy and France. The absence of an efficient swine vaccine against H1C.2.4 strains ^53^, combined with their high antigenic distance from other H1C.2 clades strains ^50^, raises concerns regarding their spread in the swine population in Europe. Interestingly H3N1 viruses were detected in Spain, Italy and Germany, which were reported in two different studies as novel subtypes ^11,54^. Phylogenetic analyses did not reveal a common origin or clear spillover patterns between countries, suggesting repeated, isolated HA/NA reassortment events. Considering that they remained sporadic over the studied decade, their fitness might be reduced compared to H3N2 or H1N1 strains. The H1A.3.3.2 lineage has become enzootic in European swine herds and has undergone major genetic/antigenic drift resulting in swine-specific clusters ^13,55^ and swIAV strains with limited cross-reaction to human seasonal viruses from this same origin ^55,56^. Additionally, H1A.3.3.2 viruses have reassorted locally with enzootic swIAV, greatly increasing the swIAV genetic diversity in Europe. For instance, a triple reassortant H3.2000.3/N2G.2/PPPPPP swIAV emerged in Danish pigs in 2014 ^42^, and was also found in Germany at that time, with a common origin according to our phylogenetic analyses. Taking into account the origin of each internal gene segment, even more novel genotypes have been identified ^11,13–15,37,38,41,42,57^ totalling 24 genotypes. According to our analyses and a recent study ^58^, the United Kingdom displayed an increase in H1A.3.3.2/N1P detection from 2020 and onward, including a clade from recent H1N1 human-to-swine spillovers dated from October 2020, which further adapted to swine. Such events were also recently reported in Denmark ^59^. This raises concerns regarding future reassortments of human-origin A/H1N1pdm09 in swine. Additionally, almost all monitored countries likely displayed H1N1 human seasonal reverse-zoonosis across the monitored period every year, which is consistent with the higher rate of human-to-swine transmissions compared to swine-to-human transmissions ^35^. France stood-out reporting the highest proportion of A/H1N1pdm09 reverse-zoonotic events compared to other countries. This is likely due to the generally low circulation of H1A.3.3.2/N1P strains in swine since 2009, and only a single swine-adapted swIAV genogroup with limited circulation compared to other swIAV lineages became established ^55^. Further comparative human IAV and swIAV phylodynamic and phylogeographic analyses are required to better understand this country-specific pattern. Frequent human-to-swine transmissions were reported for the considered decade in most analysed countries, but no avian IAV spillovers were detected in swine, suggesting that such spillovers remained rare since the emergence of the H1C/N1EA lineage.

Regarding swIAV transmissions between countries in Europe, Bayesian Markov Jumps phylogeography approach on H1C viruses revealed clear patterns of transmissions, with for instance frequent transfers from Denmark to Italy and Germany. Such transmission routes were found to be supported by live swine trades data as our Bayesian GLM phylogeographic analyses revealed that trade of live weaning pigs was responsible for swIAV transmissions between European countries. Better biosecurity measures for weaning pigs are thus probably required to limit cross-border events in Europe. Only one study, in preprint, investigated such factors driving European swIAV cross-border transmission ^60^. Among the tested factors (e.g., pig population size, organ trade, pig trade), only live-swine breeder trade significantly drove swIAV cross-border transmissions. However, that study only considered breeder live-swine trade and not weaners live-swine trade in BEAST GLM models, the sequence data were different (from 2010 to 2020) and trade data were kilogram-based instead of USD-based and from a different database. These factors limit direct comparison with our findings. The major limitation of such analyses is the availability of equivalent surveillance data in each country to avoid the introduction of biases. For the Markov Jumps analysis, several countries lacked sufficient surveillance data compared to other countries, which likely skewed certain transmission routes. For instance, few sequences were available in The Netherlands compared to Germany for the considered period, which likely shifted the phylogeographic transmission for Germany to the Netherlands rather than the converse, as found in the live swine trade data. Performing phylogeographic analyses on data from 2020 and onward should give a more accurate view of swIAV transmissions within Europe, since sequencing data availability increased over time in Europe for the considered period as shown on Figure 3. Such analyses will for instance be possible within the European Swine Influenza Network (ESFLU) COST Action which analysed swIAV diversity in Europe from October 2022 until October 2026^61^.

## Conclusion

This study provides a comprehensive overview of the genetic landscape of swIAVs in Europe between 2013 and 2022, introducing a Nextclade-based genotyping tool and harmonized nomenclature that revealed a staggering 150 genotypes circulating across the monitored countries. By standardizing classification of all eight viral segments, this framework will enhance community research and continent-wide comparison through streamlined, robust comparative analyses of swIAV genotypes. This accessible, automated genotyping approach will also allow rapid identification of zoonotic cases in human IAV samples within a One Health context. Integrating phylogeographic modelling with economic trade data identified international movement of live swine as a significant driver of swIAV spread between European countries. Repeated detection of human-to-swine spillovers across Europe highlights the constant risk of human seasonal strains seeding new diversity in swine populations, increasing the risk of reassortment and emergence of genotypes with pandemic potential. These findings underscore that harmonized, real-time surveillance is essential for effective, region-specific vaccine strategies and pre-pandemic preparedness.

## Methods

### Sample selection

Clinical samples were obtained from farmed pigs through national and regional surveillance programmes as well as focused veterinary investigations. Total RNA was extracted according to the manufacturer’s kit instructions. Chemistries used were silica membrane-based such as the QIAmp® Viral RNA Mini or QIAgen® RNeasy Mini kit (Qiagen), Nucleospin® RNA Mini Kit (Machery Nagel) or similar. Alternatively magnetic particle technology was used such as the BioSprint® One-for-all Vet (Qiagen), KingFisher™ Flex Purification System or MagMAX™ CORE nucleic acid purification (ThermoFisher Scientific), ID Gene^TM^ Mag Fast Extraction (Innovative Diagnostics) or similar kits MagNA Pure 96 DNA and Viral NA Small Volume Kit automated on the Magna Pure 96 (Roche, Switzerland). Samples were screened for the presence of viral RNA using an M-gene RT-PCR assay as described previously ^11,57,62–65^.

### Whole genome sequencing (WGS)

RNA samples were used to generate cDNA using SuperScript™ III or IV reverse transcriptase and Platinium® Taq polymerase (Invitrogen) or similar. Double-stranded cDNA was generated using sequence-independent single-primer amplification ^66^ or specific primers to generate swIAV amplicons ^65,67^. Library preparation was performed according to the manufacturers’ instructions using specific kits appropriate for the sequencing technology. The NexteraXT DNA Library Prep Kit and MiSeq Reagent Nano Kit v2 or NextSeq v2 kit were used for Illumina short-read sequencing (Illumina) on a NextSeq™ 2000 sequencer. The Ion Xpress Plus Fragment Library kit and Ion PI Hi-Q Sequencing kit (Thermo Fisher Scientific) were used for Ion Torrent sequencing (Thermo Fisher Scientific) and the Rapid Barcoding Kit and the Ligation Sequencing Kit were used for nanopore sequencing (Oxford Nanopore Technologies (ONT)). Raw sequencing reads were assembled using a custom script as described ^50,68^. Alternatively software packages were used ^57,65^ including CLC genomic workbench (Qiagen) or ONT Guppy Basecalling Software (ONT) and Geneious (Biomatters) and Bowtie 2 ^69^.

### Maximum-likelihood phylogenies with Nextstrain

The reference data set included all European swIAV data that was publicly available on GISAID EpiFlu and NCBI Virus databases within the timeframe 2009-2022 as well as data generated by the six European partner countries in the PIGIE project (Germany, Denmark, France, Italy, Spain, and the United Kingdom) and now publicly available (Supplementary Table S2). Duplicate sequences were removed using seqkit v2.7.0^70^ and further refined manually. Sequences were then aligned for each gene using Mafft v7.487 and manually trimmed to the open reading frame using AliView. Sequences with too large gaps were removed manually. Sequences metadata including collection date, country, clades for each segment, subtype, lineages, and genotypes (see below) were stored in a tab-delimited table using ad-hoc scripts. Maximum-likelihood phylogenies were generated using an augur pipeline executed within a NextStrain ^71^ cli v8.0.1 shell environment using the sequences metadata table and fasta files as input. i) Augur filter was first used to insure consistency between the sequences and the metadata. ii) Augur align performed the sequence alignment using MAFFT. iii) Augur tree with “-bb 2000 -nt 8” calculated maximum-likelihood phylogenetic trees. iv) Augur refine calculated time-scaled phylogenies with the parameters “--timetree --date-confidence --max-iter 300” using TreeTime. v)

Augur ancestral with the parameters “--inference-joint --keep-ambiguous” and augur translate were used with the A/California/07/2009 reference sequences to compute the mutations along the phylogeny branches. iv) Augur traits with the parameters “--columns country” was used to compute clades and countries memberships of the tree nodes with TreeTime mugration tests. v) Augur export v2 was used on the time-scaled phylogeny with the calculated node data to generate the auspice json. vi) An ad-hoc script using jq and sed was used to extract the infered tree leaves sampling dates, which were fed to augur frequencies with the parameters “--method kde --narrow-bandwidth 0.083 --wide-bandwidth 0.25 --proportion-wide 0.0 --min-date 2013-01-01 --max-date 2022-12-31 --pivot-interval 1” to generate the auspice frequencies json. The json files were finally uploaded on a github repository to be accessible online on the Nextstrain communities portal. Streamgraphs presented in the manuscript were built based on the Nextstrain metadata files using the streamgraph, lubridate, vroom, dplyr and htmlwidgets R packages.

### Clade definition

A swIAV sequences dataset was assembled combining the OctoFlu tool sequences and European swIAV 2009-2022 dataset. Maximum-likelihood phylogenies for each gene were generated using IQTree2 with the parameters “-bb 1000 -nt 8”. From these phylogenies, clades were defined by sharing of a common node and monophyly within swine and greater than or equal to 75% ultrafast bootstrap support based on the maximum-likelihood phylogenetic trees. Within and between clade average pairwise distance was calculated using MEGA-CC v10.2.6 ^72^ and setting the within-clade mean distance of below 5% and between-clade differences above 5% for HA and NA ^2^. These thresholds were set at 3% for the remaining gene segments ^68^. Clades defined this way were compared to the ones defined in the OctoFlu dataset and were thus confirmed. Additional clades absent from the OctoFlu dataset were defined using the analysis results and by manually curating relevant sequences by selecting distant sequences present in multiple countries with a focus on the root of the clade. These sequences as well the OctoFlu sequences were stored in a fasta file with clades name attributed to each sequence, allowing the definition of clades at the international level (including Asia and Americas).

### Nextclade classification tool building

The labelled fasta file with clade names was used alongside a metadata file describing the sequences clades in the same pipeline as the one described in the Maximum-likelihood phylogenies by Nextstrain section of this manuscript. Additionally, an automatic filter of sequences that were too short was used with “seqkit seq -u -g –m $size”, with $size being two-third of each segment maximum length, and with the “--max-iter 5” parameter for the augur refine step. A/California/07/2009 (H1N1) sequences were used as a reference and as the base root for H1, N1 and internal genes phylogenies. A/Wisconsin/67/2005 (H3N2) sequences were used as the base root and reference for the H3 and N2 phylogenies. Root sequences were further modified from the chosen reference by inferring the best fitting root through the maximum-likelihood phylogeny. The inference of the best fitting root was necessary considering the diversity of swIAVs and to limit the number of builds to 10 (H1, H3, N1, N2, PB2, PB1, PA, NP, M, NS) by avoiding the subdivision of the phylogenies at the lineage level. The obtained json phylogenetic trees were moved in a file system complying with the Nextclade community builds prerequisites containing for each segment the phylogenetic tree in json format, the root sequence, the description of CDS in gff3 format, a pathogen.json file describing the build, and a readme and a changelog file giving extra information regarding the build. They are available at: https://github.com/nextstrain/nextclade_data/tree/release/data/community/anses/swIAV. A wrapper for the Nextclade command-line was developed to easily genotype all eight segments of swIAV in a single command and store the resulting clades, subtypes, lineages and genotypes automatically in a Nextstrain-compatible metadata.txt file, which was made available at: https://github.com/gtrichard/influenza_sequences_toolbox/blob/main/bin/nextcladeGenotype. This tool was used on the previously described 2009-2022 European swIAV dataset to assign clades, subtypes, lineages and genotypes to each strain.

### Commercial exchanges data extraction and visualization

Live swine trade data were obtained from the World International Trade Solutions portal based on the UN COMTRADE database (https://wits.worldbank.org/) by extracting HS:010310 (live swine, purebred breeders) and HS:010391 (live swine, weighing less than 50 kg each) gross exports for each European country between 2013 and 2022. An annual mean was calculated between each country and the average gross exports volumes expressed in 1,000 USD were represented as Circos-like plots using an R script based on the vroom, tidyr, dplyr, data.table and circlize packages.

### Bayesian phylogeography

Two bayesian phylogenetic and phylogeographic analyses were performed using BEAST X and BEAUti v10.5.0. The dataset consisted of 1201 swIAV sequences representing the H1C lineage in countries with 10 or more sequences between 2013 and 2022, which corresponded to Denmark, Germany, Italy, France, Belgium, the Netherlands, Spain, and Austria. For each analysis, sequence evolution was modelled using the SRD06 codon-partitioned substitution model. For each partition, the Hasegawa-Kishino-Yano (HKY) substitution model was used with gamma-distributed rate variation among sites (with four rate categories). A strict molecular clock model was applied to estimate the evolutionary rates, and a coalescent constant population size tree prior was employed to model the tree-generating process. Posterior distributions of parameters were estimated using Markov chain Monte Carlo (MCMC) sampling. The MCMC chain was run for 500 million iterations, with trees and parameter values sampled every 10,000 iterations. Two analyses were conducted using these parameters.

A first analysis aimed at reconstructing the spatial diffusion of H1C swIAVs within Europe. We thus performed a discrete phylogeographic analysis on the “country” trait of each sequence based on previously described methods^73^. Markov Jumps were calculated by reconstructing the country states at all ancestors and by reconstructing state change counts along the phylogenies. The BSSVS method was additionally integrated in the run to calculate Bayes Factors using SpreaD3.

A second analysis was conducted to assess the role of live swine trades on swIAV dispersion between countries. We used a Generalized Linear Model (GLM) parameterization of the discrete trait diffusion process to simultaneously estimate the ancestral geographic states and the predictors of spatial transition rates by providing two matrices to the model. Both represented the average annual gross exports of live swine between the considered European countries, with one for live-swine weighing less than 50kg each (defined as the weaners matrix) and one for purebred breeders (defined as the breeders matrix). Values in the matrices were normalized and log transformed.

## Acknowledgements

We wish to thank research and diagnostic laboratories in Europe for generating data and making them publically available: Charlotte K. Hjulsager, Ramona Trebbien, Jesper Schak Krog from the Statens Serum Institut in Denmark ; Stéphane Gorin and Stéphane Quéguiner from the French Agency for Food, Environmental and Occupational Health & Safety in France ; Laura Baioni from the Istituto Zooprofilattico Sperimentale della Lombardia e dell’Emilia Romagna in Italy ; Ivan Domingo-Carreño and Soledad Serena from Universitat Autònoma de Barcelona in Spain. We wish to thank Alex M. P. Byrne from The Crick Institute, United Kingdom, for discussions regarding the proposed swIAV nomenclature. We with thank Tavis K. Anderson from Virus and Prion Research Unit, National Animal Disease Center, USDA-ARS, Ames, IA, USA, regarding his contributions to the establishment of the swine influenza A clades definition.

## Author contributions

G.R., B.M., P.R., H.E.E. and G.S. designed the study. G.S., H.E.E., L.E.L., C.C., T.H. and E.E.M. acquired funding. G.R. and B.M. collected data. G.R., G.S., S.H., P.R., L.E.L., A.M. and C.C. sequenced new swIAV strains and provided their sequences for the study. P.R., G.R. and B.M., with the help of G.S., L.E.L., T.H., H.E.E., M.C.M., E.M.M., S.H. and C.C. designed the whole genome swIAV nomenclature system. G.R. implemented the Nextclade swIAV genotyping tool. G.R. designed and performed data formatting, data analyses and figures. G.R., H.E.E. and G.S. wrote the paper. All authors reviewed and edited the paper.

## Ethics declaration

The authors declare no competing interests.

## Supplementary data

**Supplementary Figure S1.**
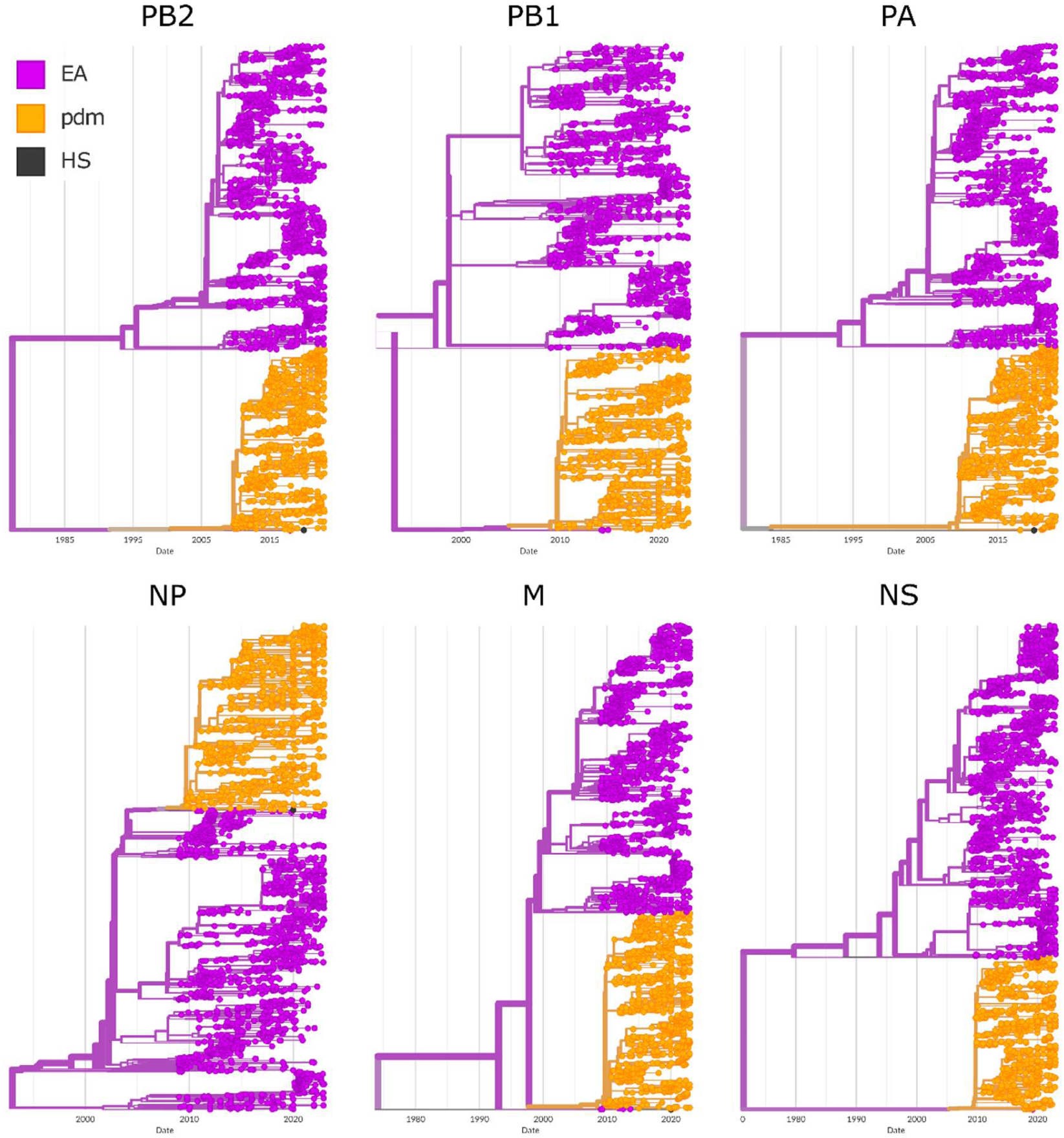
Maximum-likelihood phylogenies of European 2009-2022 swIAVs internal genes based on the novel classification. EA (purple) = Eurasian avian-like lineage, A/H1N1pdm09 (orange) = Pandemic 2009 lineage, HS (black) = human seasonal 2020 lineage (recent zoonoses).

**Supplementary Figure S2.**
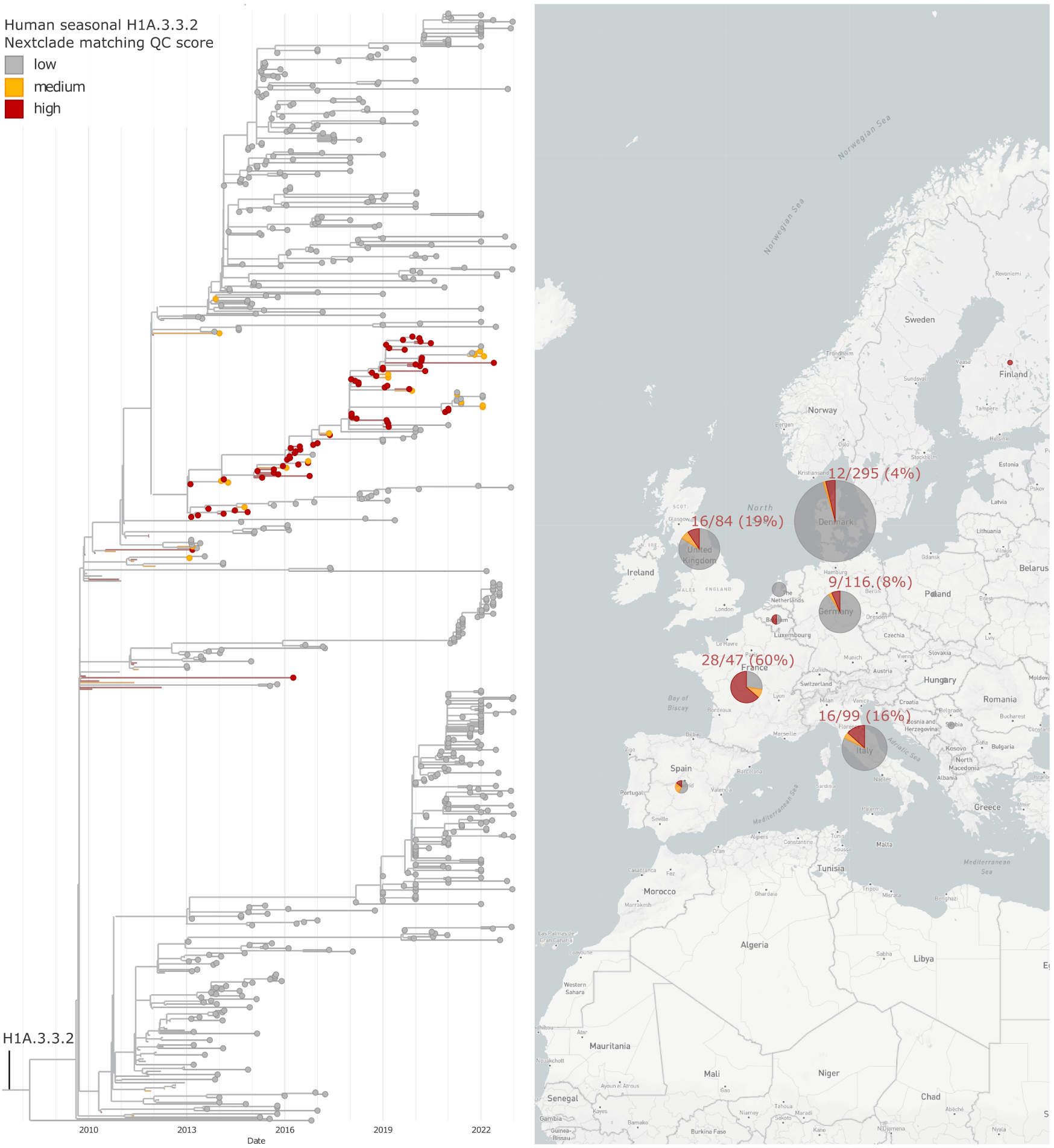
Proportion of potential reverse zoonotic transmissions per country within the H1A.3.3.2 phylogeny between 2013 and 2022. Left: swIAV H1A.3.3.2 phylogeny with sequences colored depending on the Human seasonal H1 Nextclade Quality Control score, reflecting the probability of reverse-zoonosis for swIAV sequences based on sequence match (mixed sites, private mutations, frame shifts, stop codons). Right: Map of Human seasonal H1 Nextclade Quality Control score proportions for H1A.3.3.2 swIAV sequences per country. Red: high match with H1 Human Seasonal sequences (high chance of reverse-zoonosis), Orange: medium match with H1 Human Seasonal sequences (medium chance of reverse-zoonosis or circulation and adaptation within the swine population after a reverse-zoonotic event). Grey: low match with H1 Human Seasonal sequences (swine adapted H1A.3.3.2, low probability of reverse-zoonosis). Proportions of sequences with a high match with H1 Human Seasonal sequences are displayed in red above the pie chart of each country.

**Supplementary Table S1.** SwIAV segments clades and their countries of circulation.

| Segment | Lineage name | Clade | Colloquial Name | Geographic Distribution (genotyping on all NCBI HA/NA swIAV available on the 03-05-2026) |
| --- | --- | --- | --- | --- |
| HA | Classical swine | H1A.1.1 | α-H1 | Canada, USA |
|  |  | H1A.1.1.2 |  | Canada, China, Hong Kong, Japan, South Korea, Taiwan, Thailand, UK, USA |
|  |  | H1A.1.1.3 |  | Canada, USA |
| | | H1A.2 | β-H1 | Cambodia, Canada, China, Hong Kong, Mexico, Myanmar, South Korea, Sweden, Thailand, USA |
|  |  | H1A.3.1 |  | Mexico |
| | | H1A.3.2 | γ-2-H1 | Mexico, USA |
|  |  | H1A.3.3 |  | China, Hong Kong, USA |
|  |  | H1A.3.3.1 |  | China |
|  |  | H1A.3.3.2 | pdm09 | Worldwide |
| | | H1A.3.3.3 | γ-H1 | South Korea, USA |
|  |  | H1A.3.3.3-c1 |  | USA |
|  |  | H1A.3.3.3-c2 |  | USA |
|  |  | H1A.3.3.3-c3 |  | USA |
|  |  | H1A.4 |  | USA |
| | | H1A.1.1 | α-H1 | Canada, USA |
|  |  | H1A.1.1.2 |  | Canada, China, Hong Kong, Japan, South Korea, Taiwan, Thailand, UK, USA |
|  |  | H1A.1.1.3 |  | Canada, USA |
| | | H1A.2 | β-H1 | Cambodia, Canada, China, Hong Kong, Mexico, Myanmar, South Korea, Sweden, Thailand, USA |
|  | Human seasonal (hu) | H1B.1.1 |  | Argentina, Australia, Brazil, Canada, Chile, China, France, Hong Kong, Mexico, Poland, Russia, UK, USA |
|  |  | H1B.1.1.1 |  | UK |
|  |  | H1B.1.1.2 |  | UK |
|  |  | H1B.1.1.3 |  | UK |
|  |  | H1B.1.2 |  | Spain |
|  |  | H1B.1.2.1 |  | Austria, Belgium, France, Germany, Italy, Netherlands, Spain, |
|  |  | H1B.1.2.2 |  | France, Italy |
|  |  | H1B.1.2.3 |  | France |
| | | H1B.2.1 | δ-2 | Mexico, USA |
| | | H1B.2.2.1 | δ-1a | USA |
| | | H1B.2.2.2 | δ-1b | USA |
|  |  | H1B.2.6 |  | Brazil |
|  |  | H1B.2.7 |  | Northern Ireland |
|  |  | H1B.1.1 |  | Argentina, Australia, Brazil, Canada, Chile, China, France, Hong Kong, Mexico, Poland, Russia, UK, USA |
|  |  | H1B.1.1.1 |  | UK |
|  |  | H1B.1.1.2 |  | UK |
|  | Eurasian avian (av) | H1C.1 |  | Belgium, China, Colombia, Denmark, France, Germany, Hong Kong, Indonesia, Ireland, Mexico, Netherlands, South Korea, Spain, Taiwan, UK, USA |
|  |  | H1C.2 |  | Denmark, Germany |
|  |  | H1C.2.1 |  | Austria, Belgium, China, Croatia, Denmark, France, Germany, Hong Kong, Hungary, Italy, Netherlands, Poland, Russia, Serbia, Spain, Sweden, Switzerland |
|  |  | H1C.2.2 |  | Austria, Belgium, China, Czech Republic, Denmark, France, Germany, Hong Kong, Italy, Netherlands, Poland, Spain, Thailand, UK |
|  |  | H1C.2.3 |  | Cambodia, China, Hong Kong, South Korea, Viet Nam |
|  |  | H1C.2.4 |  | Croatia, Denmark, Finland, Germany, Italy, Poland, Spain, Thailand |
|  |  | H1C.2.4.1 |  | Denmark, France, Germany, Italy, Netherlands, Poland, Spain, Ukraine |
|  |  | H1C.2.4.2 |  | Denmark, France, Germany, Italy, Spain |
|  |  | H1C.2.4.3 |  | Denmark, France, Italy |
|  |  | H1C.2.5 |  | Denmark, Germany, Italy, Poland, Sweden |
|  |  | H1C.2.6 |  | Italy, Spain |
|  | Human seasonal swine-adapted | H3.1970.1 |  | Austria, Belgium, Cambodia, China, Denmark, France, Germany, Hungary, Italy, Mexico, Myanmar, Netherlands, Singapore, South Korea, Spain, Switzerland, Taiwan, Thailand, USA |
|  |  | H3.1990.1 |  | Argentina, Australia, Brazil, Canada, Chile, China, Germany, Italy, Japan, Kazakhstan, Mexico, Russia, South Korea, Taiwan, Thailand, UK, USA |
|  |  | H3.1990.4 |  | Canada, Mexico, South Korea, USA |
|  |  | H3.1990.4.a |  | South Korea, USA |
|  |  | H3.1990.4.b1 |  | Canada, USA |
|  |  | H3.1990.4.b2 |  | Canada, USA |
|  |  | H3.1990.4.d |  | USA |
|  |  | H3.1990.4.e |  | USA |
|  |  | H3.1990.4.f |  | USA |
|  |  | H3.1990.4.g |  | Canada, South Korea, USA, Viet Nam |
|  |  | H3.1990.4.h |  | USA |
|  |  | H3.1990.4.i |  | Canada, USA |
|  |  | H3.2000.3 |  | Denmark, Germany, Spain, |
|  |  | H3.2010.1 |  | Cambodia, China, Guatemala, Hong Kong, Sri Lanka, USA |
|  |  | H3.2010.2 |  | India, USA |
|  |  | H3.2020.1 |  | USA |
|  |  | H3.2020.2 |  | USA |
|  |  | H3.Other-Human-2010 |  | Denmark, Guatemala, Hong Kong, Italy, USA, Zambia |
|  |  | H3.Other-Human-2020 |  | Chile, Kazakhstan, USA |
| NA | Classical swine | N1.C.1 |  | Brazil, Canada, China, Hong Kong, Japan, Mexico, Singapore, South Korea, Sweden, Switzerland, Taiwan, UK, USA |
|  |  | N1.C.1.1 |  | Argentina, Brazil, Canada, China, Hong Kong, Russia, Taiwan, USA |
|  |  | N1.C.1.2 |  | Canada, USA |
|  |  | N1.C.2 |  | Mexico, South Korea, USA |
|  |  | N1.C.2.1 |  | USA |
|  |  | N1.C.3 |  | Brazil, Canada, China, Hong Kong, Mexico, South Korea, Taiwan, UK, USA |
|  |  | N1.C.3.1 |  | USA |
|  |  | N1.C.3.2 |  | USA |
|  | Eurasian avian (av) | N1EA.1 |  | Austria, Belgium, Croatia, Denmark, France, Germany, Italy, Netherlands, Poland, Russia, Serbia, Spain, Switzerland, Thailand, UK |
|  |  | N1EA.2 |  | Austria, Belgium, France, Germany, Hungary, Italy, Netherlands, Poland, Portugal, Russia, Spain |
|  |  | N1EA.3 |  | Croatia, Denmark, Finland, Germany, Italy, Poland, Spain, Sweden |
|  |  | N1EA.4 |  | Belgium, Cambodia, China, Croatia, Czech Republic, Denmark, France, Germany, Hong Kong, Indonesia, Ireland, Italy, Mexico, Netherlands, Poland, South Korea, Spain, Thailand, UK, USA, Viet Nam |
|  | Pandemic 2009 (pdm) | N1P | pdm09 | Worldwide |
|  | Human seasonal swine-adapted | N2.1998A |  | USA |
|  |  | N2.1998B |  | USA |
|  |  | N2.2002 |  | Argentina, Australia, Brazil, Cambodia, Canada, China, France, Japan, Mexico, Russia, South Korea, USA, Viet Nam |
|  |  | N2.2002A |  | South Korea, USA |
|  |  | N2.2002B |  | Canada, USA |
|  |  | N2.2016 |  | Brazil, Cambodia, China, Denmark, Guatemala, Hong Kong, Sri Lanka, USA, Viet Nam |
|  | European Human seasonal swine-adapted (derived from 1990s human viruses) | N2EU |  | Germany |
|  |  | N2EU.3 |  | Spain |
|  |  | N2EU.4 |  | Italy |
|  |  | N2EU.5 |  | Denmark |
|  |  | N2EU.6 |  | Denmark |
|  |  | N2EU.7 |  | Northern Ireland |
|  | Gent/1984 | N2G.1 |  | Austria, Belgium, Denmark, France, Germany, Hong Kong, Hungary, Italy, Netherlands, Poland, Spain |
|  |  | N2G.2 |  | Belgium, Denmark, France, Germany, Italy, Netherlands, Poland, Spain, Sweden, Ukraine |
|  | Human seasonal (2020) | N2HS |  | Chile, Denmark, India, Kazakhstan, USA, Zambia |
|  | Texas/1985 | N2LAIV98 |  | Australia, Brazil, Cambodia, Canada, Chile, China, Germany, Hong Kong, Japan, Mexico, Myanmar, South Korea, Thailand, South Korea, Thailand, USA, Viet Nam |
|  | Scotland/1994 | N2S |  | Australia, Austria, Belgium, China, France, Germany, Hong Kong, Italy, Japan, Mexico, Netherlands, Poland, South Korea, Spain, Taiwan, Thailand, UK, USA |
| PB2,<br>PB1,<br>PA, NP,<br>M, NS | Human<br>seasonal<br>(2020) | HS (H) |  |  |
|  | Eurasian<br>avian | EA ( E) |  |  |
|  | 2009<br>pandemic | pdm (P) | pdm09 |  |
|  | Triple<br>Reassortant<br>Genotype | TRIG (T) |  |  |
|  | Eurasian<br>avian spillover | spillover (S) |  |  |
|  | Texas/1985 | LAIV (L) |  |  |

**Supplementary Table S2.**
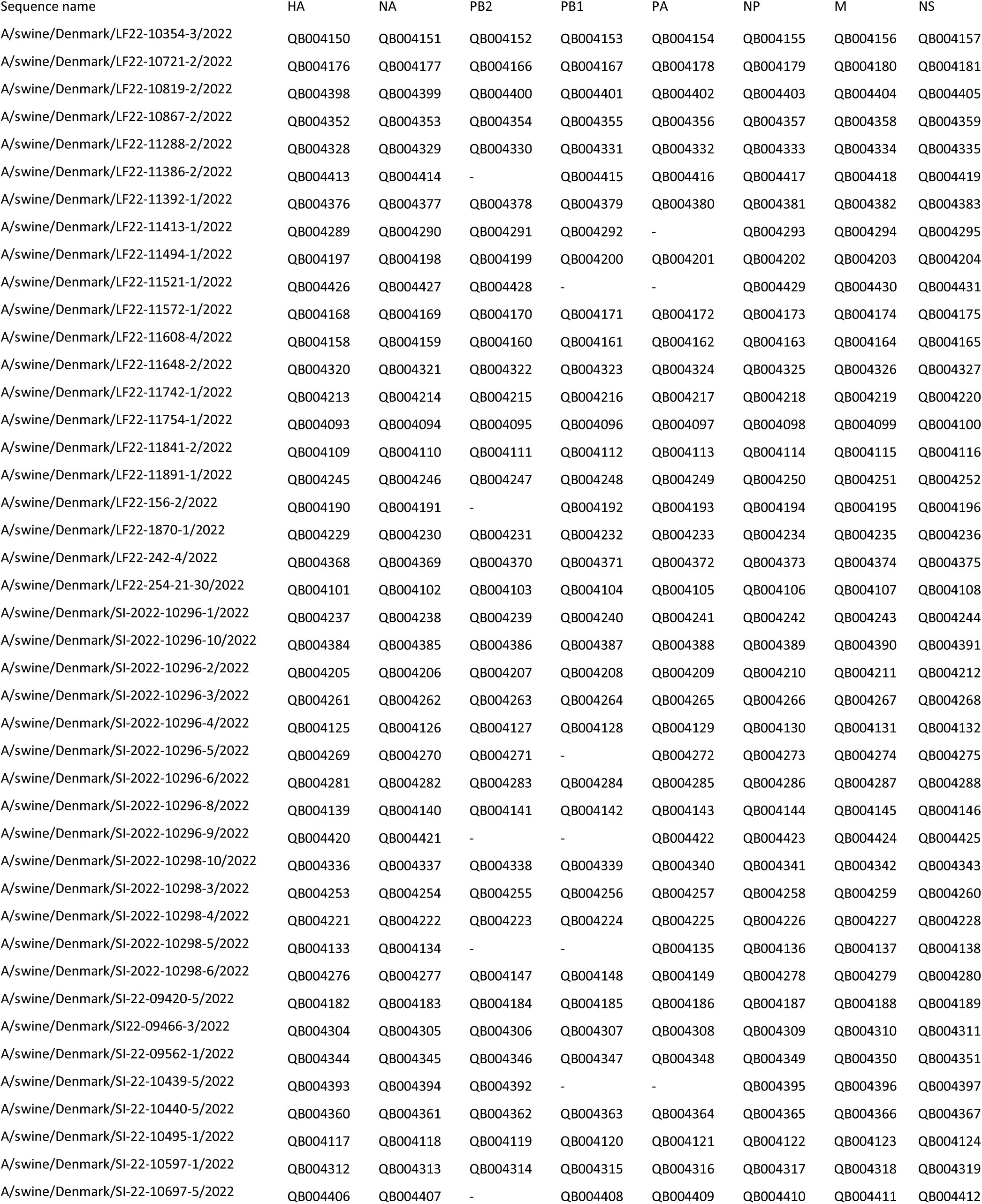
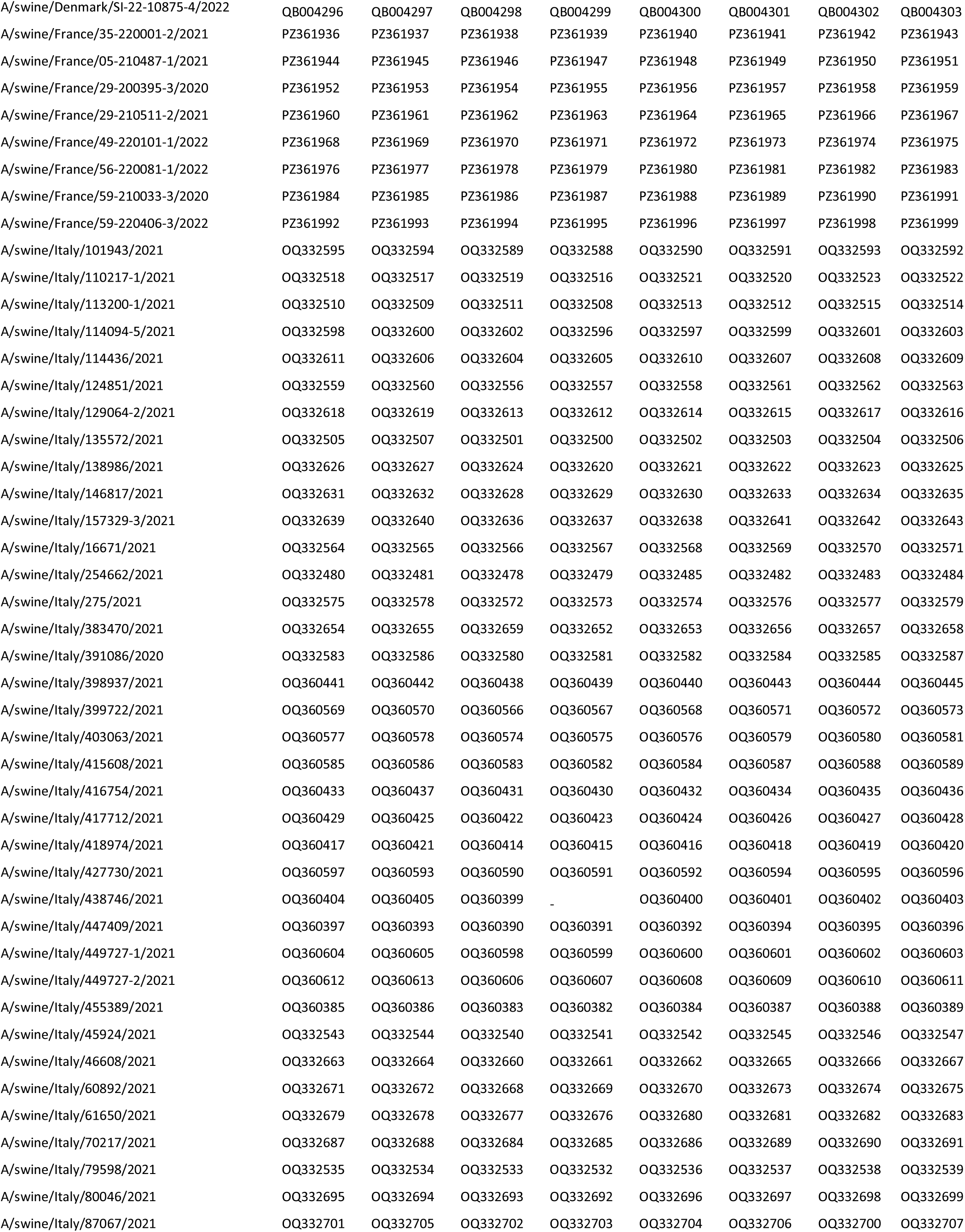
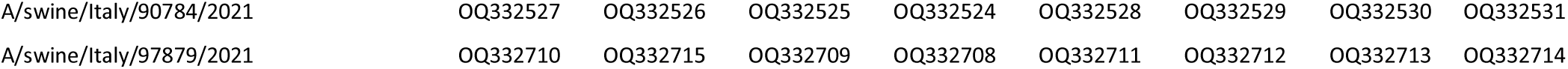
Sequenced strain names and related GenBank IDs.

**Supplementary Table S3.** BEAST runs quality control statistics used to generate the H1C phylogeographic Markov Jumps analyses in Figure 5A, 5B, and 5C.

| BEAST run QC Statistics and information | Run 1 ESS | Run2 ESS | Run 3 ESS | Combined ESS |
| --- | --- | --- | --- | --- |
| Precision | double | double | double | double |
| Sequences | 1201 | 1201 | 1201 | 1201 |
| Sites | 1584 | 1584 | 1584 | 1584 |
| states | 178 940 000 | 338 060 000 | 175 150 000 | 622 950 000 |
| burn-in | 17 893 000 | 33 805 000 | 17 515 000 | - |
| joint | 304 | 635 | 422 | 435 |
| prior | 114 | 166 | 179 | 402 |
| likelihood | 1506 | 3652 | 1161 | 279 |
| treeModel.rootHeight | 1255 | 302 | 1663 | 887 |
| age(root) | 1255 | 302 | 1663 | 887 |
| treeLength | 84 | 129 | 131 | 308 |
| constant.popSize | 339 | 686 | 500 | 1362 |
| default.clock.rate | 212 | 511 | 353 | 493 |
| country.clock.rate | 3982 | 7816 | 4723 | 15111 |
| default.meanRate | 212 | 511 | 353 | 493 |
| country.meanRate | 3982 | 7816 | 4723 | 15111 |
| default.treeLikelihood | 1326 | 3004 | 1227 | 282 |
| coalescent | 96 | 138 | 145 | 332 |
| c_country.count | 1597 | 3703 | 2141 | 7147 |
| c_allTransitions | 1602 | 3695 | 1930 | 7260 |

**Supplementary Table S4.** Quality control statistics and results obtained from a BEAST analysis of H1C swIAV sequences from 2013 to 2022 with the weaners (HS:010391) and purebred breeders live-swine (HS:010310) gross exports exchanges between European countries set within a Generalized Linear Model (log10 normalized).

| BEAST Run Description | Run 1 | Run 2 | Combined |
| --- | --- | --- | --- |
| Precision | double | double | double |
| Number of sequences | 1201 | 1201 | 1201 |
| Sites | 1584 | 1584 | 1584 |
| States | 286 730 000 | 285 760 000 | 515 250 000 |
| Burn-in | 28 673 000 | 28 576 000 | - |

| QC Statistics | Mean | ESS | Mean | ESS | Mean | ESS |
| --- | --- | --- | --- | --- | --- | --- |
| joint | -1.30E+05 | 836 | -1.30E+05 | 730 | -1.30E+05 | 1648 |
| prior | -8340.96 | 308 | -8340.76 | 334 | -8340.863 | 646 |
| likelihood | -1.22E+05 | 5537 | -1.22E+05 | 3135 | -1.22E+05 | 7096 |
| rootHeight | 31.409 | 62 | 31.285 | 33 | 31.347 | 33 |
| age(root) | 1991.586 | 62 | 1991.709 | 33 | 1991.647 | 33 |
| treeLength | 3871.915 | 253 | 3871.223 | 278 | 3871.57 | 530 |
| constant.popSize | 356.075 | 1032 | 356.089 | 1069 | 356.082 | 2093 |
| default.clock.rate | 3.55E-03 | 604 | 3.55E-03 | 586 | 3.55E-03 | 1184 |
| country.clock.rate | 1.66E-02 | 8851 | 1.66E-02 | 8822 | 1.66E-02 | 17700 |
| default.meanRate | 3.55E-03 | 604 | 3.55E-03 | 586 | 3.55E-03 | 1184 |
| country.meanRate | 1.66E-02 | 8851 | 1.66E-02 | 8822 | 1.66E-02 | 17700 |
| default.treeLikelihood | -1.21E+05 | 5522 | -1.21E+05 | 1437 | -1.21E+05 | 4890 |
| country.treeLikelihood | -715.345 | 7253 | -715.311 | 5652 | -715.328 | 12384 |
| default.branchRates | 0.00E+00 | - | 0.00E+00 | - | 0.00E+00 | - |
| country.branchRates | 0.00E+00 | - | 0.00E+00 | - | 0.00E+00 | - |
| coalescent | -8250.08 | 289 | -8249.91 | 318 | -8249.997 | 609 |

| GLM Results | Mean | ESS | Mean | ESS | Mean | ESS |
| --- | --- | --- | --- | --- | --- | --- |
| country.includedPredictors | 1.066 | 24964 | 1.068 | 25063 | 1.067 | 50025 |
| country.coefficients_breeders | 2.99E-03 | 24864 | 2.76E-03 | 25399 | 2.87E-03 | 50951 |
| country.coefficients_weaners | 1.689 | 20789 | 1.686 | 19880 | 1.687 | 40610 |
| country.coefIndicators_breeders | 6.65E-02 | 24964 | 6.77E-02 | 25063 | 6.71E-02 | 50025 |
| country.coefIndicators_weaners | 1 | - | 1 | - | 1 | - |
| country.coefficientsTimesIndicators_breeders | -3.80E-03 | 25807 | -3.65E-03 | 24014 | -3.73E-03 | 51527 |
| country.coefficientsTimesIndicators_weaners | 1.689 | 20789 | 1.686 | 19880 | 1.687 | 40610 |

